# Spatial organization of plant defense at the infection front

**DOI:** 10.64898/2026.04.01.715938

**Authors:** Xiaopeng Li, Yi-Chang Sung, Gitta Coaker, Jie Zhu

**Affiliations:** Division of Biological Sciences, Interdisciplinary Plant Group, University of Missouri-Columbia, Columbia, MO, 65211, USA; Department of Plant Pathology, University of California, Davis, Davis, CA, 95616, USA

**Keywords:** Plant immunity, Spatial organization, single-cell resolution

## Abstract

Plants lack specialized immune cells and instead rely on coordinated cellular responses to restrict pathogen invasion while preserving tissue integrity. How plant immunity spatially organizes these responses remains unclear. Using live-cell reporters in *Arabidopsis* infected with *Pseudomonas syringae*, we show that effector-triggered immunity, superimposed on pattern-triggered immunity, establishes a sustained yet spatially confined immune architecture at the infection front. Defense activation persists for days but remains restricted to a narrow ring of cells surrounding viable bacterial microcolonies. Over time, immune activity spreads to adjacent layers, forming a coordinated, multilayered defense zone. This zonation extends beyond transcriptional activation to polarized callose deposition at pathogen-facing cell walls, reinforcing a localized containment boundary that limits pathogen spread. Consistent with previous single-reporter studies, simultaneous visualization of salicylic acid (SA) and jasmonic acid (JA) biosynthesis and response markers reveals a radial hormone gradient, with SA-enriched cells proximal to bacterial colonies and JA-enriched cells in surrounding regions. Individual cells predominantly activate one pathway, indicating that SA–JA antagonism is resolved through spatial compartmentalization across neighboring cells. Together, these findings establish immune zonation as a strategy for robust pathogen containment while minimizing collateral tissue damage.

**Significance statement:** Plants lack specialized immune cells and must coordinate defense responses across tissues to contain infection. Although plant immune responses are known to be locally activated, how they are organized across cells to form a stable containment boundary has remained unclear. Using live imaging of fluorescent bacterial microcolonies, we show that plant immunity forms a spatially organized defense zone surrounding viable infection sites. Within this zone, distinct defense activities are arranged in a radial gradient, with salicylic acid–associated responses enriched in cells near the pathogen and jasmonic acid–associated responses in surrounding cells. Rather than being resolved solely within individual cells, antagonistic hormone pathways are separated across neighboring cells. This spatial organization reveals immune zonation as a strategy for confining pathogen spread while preserving tissue integrity.

## Introduction

Plants rely on innate immune systems to detect and restrict pathogen invasion while maintaining tissue integrity and growth (1–3). Upon recognition of pathogen-derived molecules and proteins by cell-surface pattern-recognition receptors (PRRs) or intracellular effectors by nucleotide-binding leucine-rich repeat (NLR) receptors, plants activate pattern-triggered immunity (PTI) and effector-triggered immunity (ETI), respectively (4). NLR activation results in strong defense and are frequently associated with hypersensitive cell death (4).

Rather than operating as independent layers, PTI and ETI mutually potentiate one another, with cell-surface and intracellular receptors cooperatively amplifying immune outputs (5, 6). This activation results in a suite of responses, including Ca²⁺ influx, transcriptional reprogramming, phytohormone production, reactive oxygen species (ROS) accumulation, and callose deposition. Although these responses effectively restrict pathogen proliferation, excessive or misregulated immune activation can compromise plant development and fitness (3, 7–10). Unlike animals, plants lack specialized mobile immune cells and instead rely on coordinated cellular responses across tissues to contain infection (2, 11, 12). This decentralized immune system necessitates precise spatial control, ensuring that defense activation is confined to regions surrounding infection sites while minimizing detrimental effects on uninfected tissues (13). Although immune responses are known to be locally restricted, how ETI reorganizes tissue-level defense outputs across neighboring cells to form a stable containment architecture remains unresolved.

Emerging evidence indicates that plant immune responses are spatially organized rather than uniformly activated. Single-cell and spatial transcriptomics have revealed immune-enriched cell populations surrounding infection sites (14–17). Fluorescent reporter imaging has similarly shown confined activation of immune marker genes such as *flagellin-induced receptor-like kinase 1* (*FRK1*) in cells adjacent to bacterial colonies (14, 18–20). These studies demonstrated spatial and cellular variation in defense using single-reporter imaging or single-cell transcriptomic analyses.

Plant immune responses are coordinated by interwoven phytohormone signaling networks, including salicylic acid (SA), jasmonic acid (JA), ethylene, and additional regulatory sectors. Genetic analyses of higher-order mutants demonstrated that disease resistance emerges from combinatorial and compensatory interactions among these pathways rather than from linear signaling modules (21–23). Among these networks, SA and JA represent a prominent antagonistic regulatory axis. During infection by hemibiotrophic pathogens such as *Pseudomonas syringae*, both SA- and JA- associated pathways are activated (24–26). Although SA–JA antagonism has been extensively characterized at the molecular level, recent work has revealed roles for jasmonate signaling during ETI and systemic immune activation (27). Advances in biosensor development have enabled high-resolution visualization of SA dynamics during pathogen infection, revealing spatial heterogeneity in SA accumulation associated with virulent and avirulent bacterial strains (28). In parallel, single-reporter analyses have suggested spatial separation of SA- and JA-associated responses at infection sites (19, 29, 30).

Here, we combine live-cell reporters with pathogen localization to define the spatial organization of immune activation during ETI in the context of PTI. We find that immune activation forms confined yet sustained defense zones surrounding viable bacterial colonies. These zones are reinforced by polarized defense outputs, including callose deposition at pathogen-facing plant cells. Simultaneous visualization of SA and JA biosynthesis and response reveal a radial hormone gradient, with SA-enriched cells proximal to bacterial colonies and JA-enriched cells in surrounding regions. Individual cells predominantly activate one pathway, indicating that SA–JA antagonism is resolved through spatial partitioning across neighboring cells. Together, these findings define immune zonation as a spatial strategy that enables pathogen containment while limiting collateral tissue damage.

## Results

### ETI-potentiated immunity establishes a stable, localized immune zone at bacterial colonization sites

To investigate how immune-activated cells are organized in space and time following PTI potentiated by ETI (ETI-potentiated immunity), we monitored expression of the defense-associated gene *FRK1* in *Arabidopsis* using the reporter line *pFRK1::NLS–3xmVenus* after flood inoculation with *Pseudomonas syringae*. *FRK1* encodes a receptor-like kinase that serves as a well-established marker of PTI activation (31, 32), and its expression is strongly potentiated by ETI (5, 6). The reporter has been widely used to monitor PTI activation in response to MAMP treatment and pathogen infection (14, 18). In our previous work, we showed that *FRK1* is strongly induced in cells adjacent to bacterial colonization sites during early virulent infection with *P. syringae* pv. *tomato* (*Pst*) DC3000 but is subsequently suppressed at later time points (14).

Infection of two-week-old *Arabidopsis* with *Pst* DC3000 carrying the effector AvrRpm1, recognized by the CC-NLR Resistance to *Pseudomonas syringae* pv. *maculicola* 1(RPM1) (33, 34), induced a distinct spatiotemporal pattern of *FRK1* expression. At 24 hours post-inoculation (hpi), *FRK1* expression was strongly induced, resembling early responses observed during virulent *Pst* DC3000 infection (Fig. 1A–B). However, unlike the virulent interaction, *FRK1* expression remained robustly elevated at 72 hpi, indicating sustained immune activation upon ETI induction. Despite this prolonged activation, *FRK1* expression was spatially confined to cells surrounding individual bacterial microcolonies rather than spreading uniformly across infected tissue (Fig. 1C). Line-scan analysis confirmed that the *FRK1* signal was highest in cells directly adjacent to bacterial colonies and declined sharply with increasing distance (Fig. 1C). These discrete clusters of *FRK1*-expressing cells, hereafter referred to as “bundles,” were associated with viable, fluorescent bacterial microcolonies at 24, 72, and 140 hpi (Fig. 1D, S1A-C). In four-week-old *Arabidopsis* leaves, *FRK1* expression remained spatially restricted following both spray inoculation and syringe infiltration (Fig. S1D). Notably, high-inoculum infiltration with *Pst* DC3000 (AvrRpm1; OD600 = 0.2) resulted in low or undetectable *FRK1* expression within the infiltrated area, while elevated expression was consistently observed at the infiltration border (Fig. S1E). This pattern closely resembles previously reported spatial restriction of the defense marker *Pathogenesis-Related protein 1 (PR1)* under similar conditions (19, 29). Collectively, these data indicate that sustained immune activation surrounds the sites of active bacterial colonization, establishing spatially restricted defense zones at infection sites.

**Figure 1.**
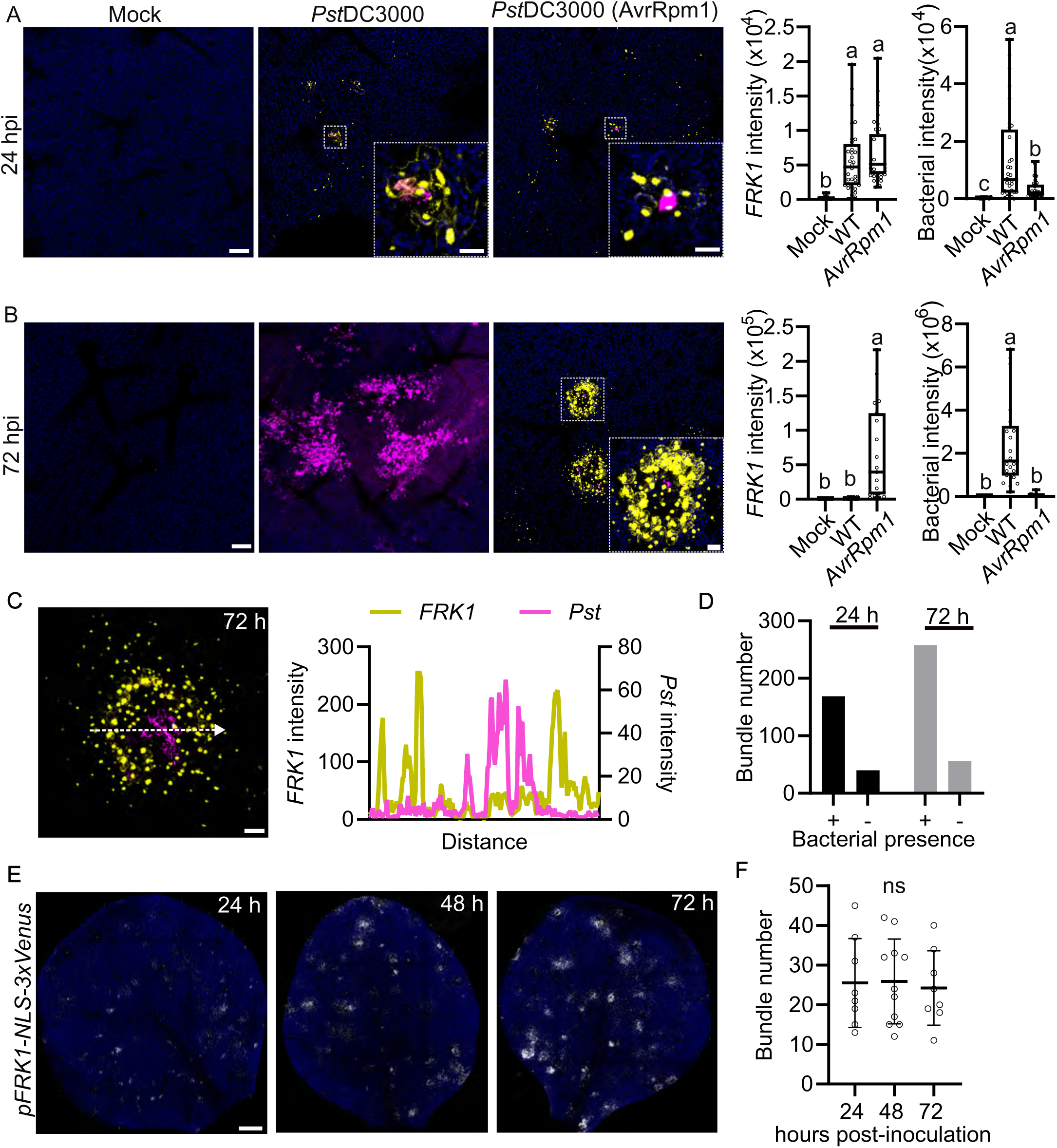
ETI-potentiated immunity establishes sustained spatial immune zones that limit infection. Two-week-old *Arabidopsis* seedlings expressing the reporter *pFRK1::NLS-3xmVenus* were flood-inoculated with *Pseudomonas syringae* pv. tomato (*Pst*) DC3000 strains expressing mCherry to visualize immune activation and bacterial colonization. Images are maximum projections of confocal z-stacks. **(A)** Spatial pattern of immune activation during virulent and avirulent infection. Seedlings were infected with virulent *Pst* DC3000 or avirulent *Pst* DC3000 (AvrRpm1). **Left:** *FRK1* expression (yellow) and bacterial signal (magenta) were imaged at 24 h post inoculation (hpi). Blue = chlorophyll autofluorescence, scale bar, 100 μm, inset bar, 25 μm. **Right:** Quantitative analysis of fluorescence signal intensities of mVenus and mCherry. Total fluorescence intensity density per colony was measured, and boxplots show median with minimum and maximum values (n ≥ 6 images from at least 3 plants). Different letters indicate statistically significant differences (*p* < 0.0001, ANOVA with Tukey test). **(B)** Infection as in (A), imaged at 72 hpi. **(C)** Zoomed view of a representative infection site at 72 hpi showing a bacterial microcolony surrounded by *FRK1*-expressing cells. Line-scan analysis across the indicated region shows the spatial distribution of *FRK1* and bacterial fluorescence signals. **(D)** Association between *FRK1* expression bundles and bacterial colonization. *FRK1* expression forms discrete clusters of responding cells (“bundles”). The presence or absence of bacterial colonies at the center of *FRK1* bundles was quantified at 24 and 72 hpi following infection with *Pst* DC3000 (AvrRpm1). **(E)** Whole-leaf imaging of *FRK1* reporter activity following infection with *Pst* DC3000 (AvrRpm1). *FRK1* expression is shown in white and chlorophyll autofluorescence in blue. Scale bar, 0.5 mm. **(F)** Quantification of the number of *FRK1* expression bundles as shown in (E). No statistically significant differences were detected (ANOVA with Tukey test, n ≥ 6 images from at least 3 plants).

To determine whether spatially restricted defense is a common feature of NLR mediated resistance, we examined *FRK1* expression following infection with *Pst* DC3000 carrying the effector AvrRpt2 or AvrRps4, recognized by the CC-NLR Resistance to *P. Syringae* 2 (RPS2) and the TIR-NLR Resistance to *P. Syringae* 4 (RPS4), respectively (34–37). In both interactions, *FRK1* expression was similarly confined to cells proximal to bacterial colonies (Fig. S1F). Spatially restricted expression was also observed for two additional defense reporters: *CBP60g*, a transcription factor associated with SA-related defense (14, 38, 39), and *LipoP1*, a putative membrane lipoprotein enriched in immune-activated cell populations (Fig. S1G) (14). Together, these results indicate that spatially confined defense gene expression is a conserved feature of NLR-mediated resistance.

### ETI activation suppresses subsequent bacterial invasion

Next, we sought to determine if ETI-potentiated immunity limits the spread of new infection events across the leaf. Bacterial colonies were relatively small and more difficult to visualize directly, whereas immune marker bundles were readily detectable. Bacterial colonies were detected in 80.9% (169/209) of *FRK1* bundles at 24 hpi and 82.2% (258/314) at 72 hpi (Fig. 1D). Therefore, we used *FRK1* bundle number as a proxy for colonization sites in whole-leaf imaging. Two-week-old *Arabidopsis* seedlings expressing fluorescent reporters were flood-inoculated with *Pst* DC3000 (AvrRpm1) at a low concentration (OD600 = 0.0005). Imaging of entire leaves revealed bundle number remained stable across 24, 48, and 72 hpi for both *FRK1* and *CBP60g* reporters (∼25 bundles per leaf at all time points; Fig. 1E–F; Fig. S2A–B), indicating that ETI limits the establishment of new infection sites rather than eliminating existing ones. In contrast, bundle size increased significantly over time for both markers (Fig. S2C–D), consistent with constrained local expansion of existing colonies. *CBP60g* bundles were generally larger than *FRK1* bundles (Fig. S2A-D). Bacterial population assays confirmed overall growth suppression during NLR activation (Fig. S2E).

### Immune activation propagates outward from the infection front in a cell layer dependent manner

The sustained *FRK1* expression observed at bacterial colonization sites raised the question of how immune activation is spatially organized across cell layers over time. To address this, we examined *FRK1* expression at single-cell resolution relative to bacterial microcolonies at sequential time points following flood inoculation with *Pst* DC3000 (AvrRpm1). At 24 hpi, *FRK1* expression was highest in cells of the first layer directly contacting bacterial colonies above substomatal cavities (Fig. 2A and SI video 1). By 36 hpi, *FRK1* expression in first-layer cells remained elevated, while second-layer cells showed a significant increase in reporter signal (Fig. 2B and SI video 2), suggesting that immune activation had begun to propagate outward from the infection front.

**Figure 2.**
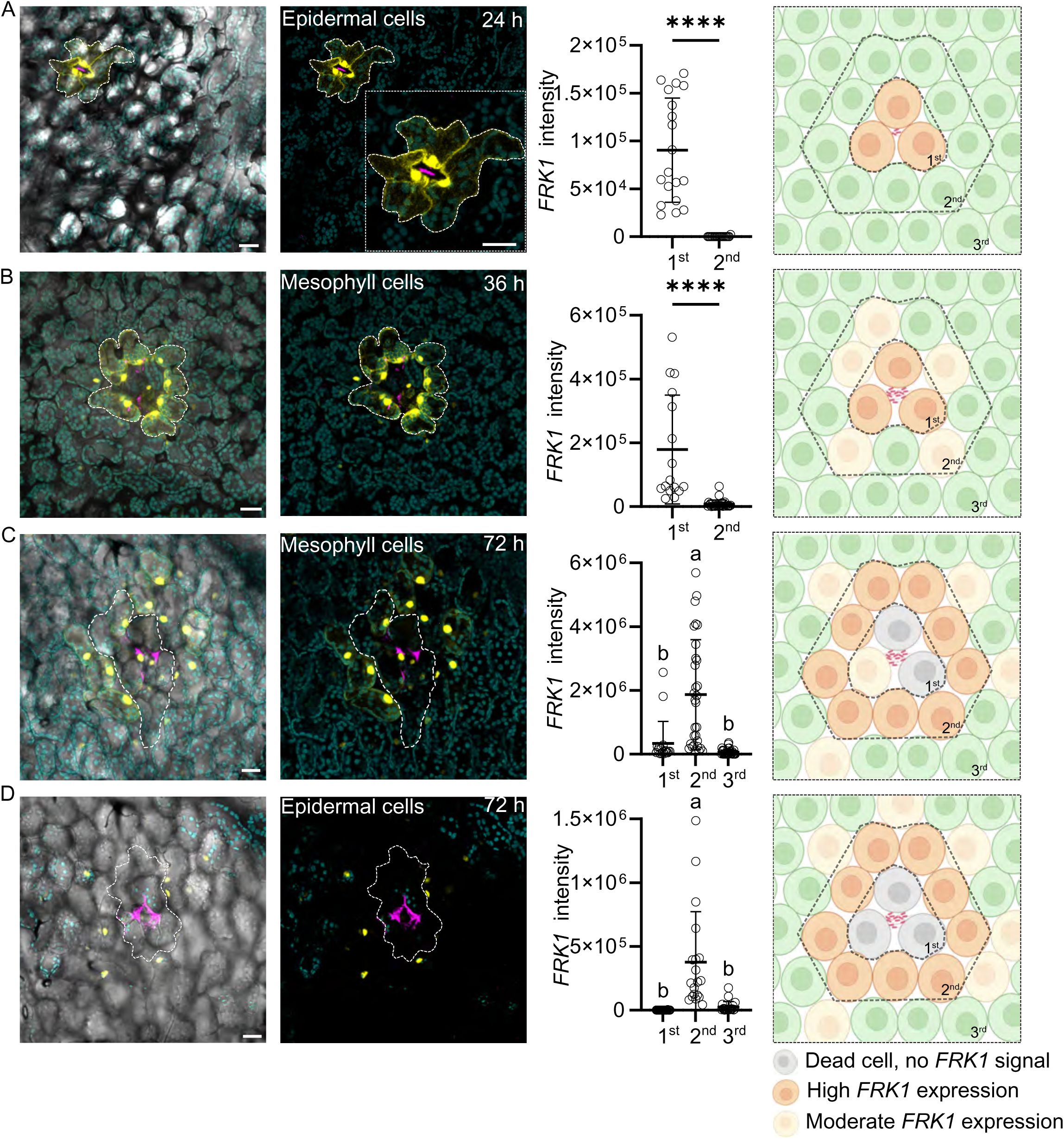
Spatiotemporal redistribution of *FRK1* expression at the infection front. Two-week-old *Arabidopsis* seedlings expressing *pFRK1::NLS–3xmVenus* were flood-inoculated with *Pst* DC3000 strains expressing mCherry or mock treated with 5 mM MgCl₂. Images are maximum projections of confocal z-stacks. Chlorophyll autofluorescence is shown in cyan. **(A-B)** *FRK1* expression in the first cell layer directly contacting bacterial colonies at 24 and 36 hpi. Left panels show *FRK1* signal at infection sites, dotted line indicates the first cell layer in direct contact with bacteria. Middle panels show quantification of *FRK1* expression in cells of the first and second layers relative to bacterial colonization. Scale bar, 25 μm. Total fluorescence intensity per cell was measured, and scatter plots show the median ± SD (n ≥ 4 infection sites), significance was determined by two-tailed Student’s t-test. Right panels are models illustrating quantitative data. **(C-D)** *FRK1* expression in second-layer cells at 72 hpi. Experiments were performed as described in (A-B). Different letters indicate statistically significant differences (*p* < 0.0001, ANOVA with Tukey test).

By 72 hpi, this spatial redistribution was more pronounced. *FRK1* expression in first-layer cells was reduced relative to the 24 hpi peak, whereas second-layer cells exhibited robust and significantly increased *FRK1* signal (Fig. 2C–D and SI video 3). This pattern indicates a temporal shift in the peak of immune activation from the innermost cell layer toward more distal layers, consistent with progressive propagation of defense signaling outward from the colonization site. Such outward propagation may reflect the spread of immune signals from the infection center, including ROS, damage, and phytocytokines (13). Together, these data reveal that ETI-potentiated immunity does not activate uniformly across cells but instead spreads in a cell-layer-dependent manner, building a multilayered defense zone over time.

### Callose deposition is spatially restricted and directionally biased toward infection sites

Callose deposition at the cell wall is a hallmark immune response that physically reinforces cell boundaries and can restrict pathogen spread in some contexts (40–42). To determine whether callose deposition is spatially organized relative to infection sites during ETI-potentiated immunity, we visualized callose accumulation by alanine blue staining in *Arabidopsis* seedlings infected with *Pst* DC3000 (AvrRpm1). Bacterial presence was identified through dead epidermal cells visualization and bacterial colonization at stomata in epidermal tissues (Fig 3A, 3E). Callose foci were detected in both epidermal and mesophyll cells proximal to bacterial microcolonies at 48 hpi (Fig. 3A–B). Foci were significantly larger within 20 µm of infection sites and decreased in size farther away from infection sites (Fig. 3C). Notably, callose accumulated preferentially in cells within approximately five cell layers of infection sites (proximal), whereas cells in distal regions showed significantly fewer and smaller foci (Fig. 3D). These data indicate that callose deposition, like *FRK1* transcriptional activation, is spatially confined to cells at the infection front rather than uniformly distributed across infected tissue.

**Figure 3.**
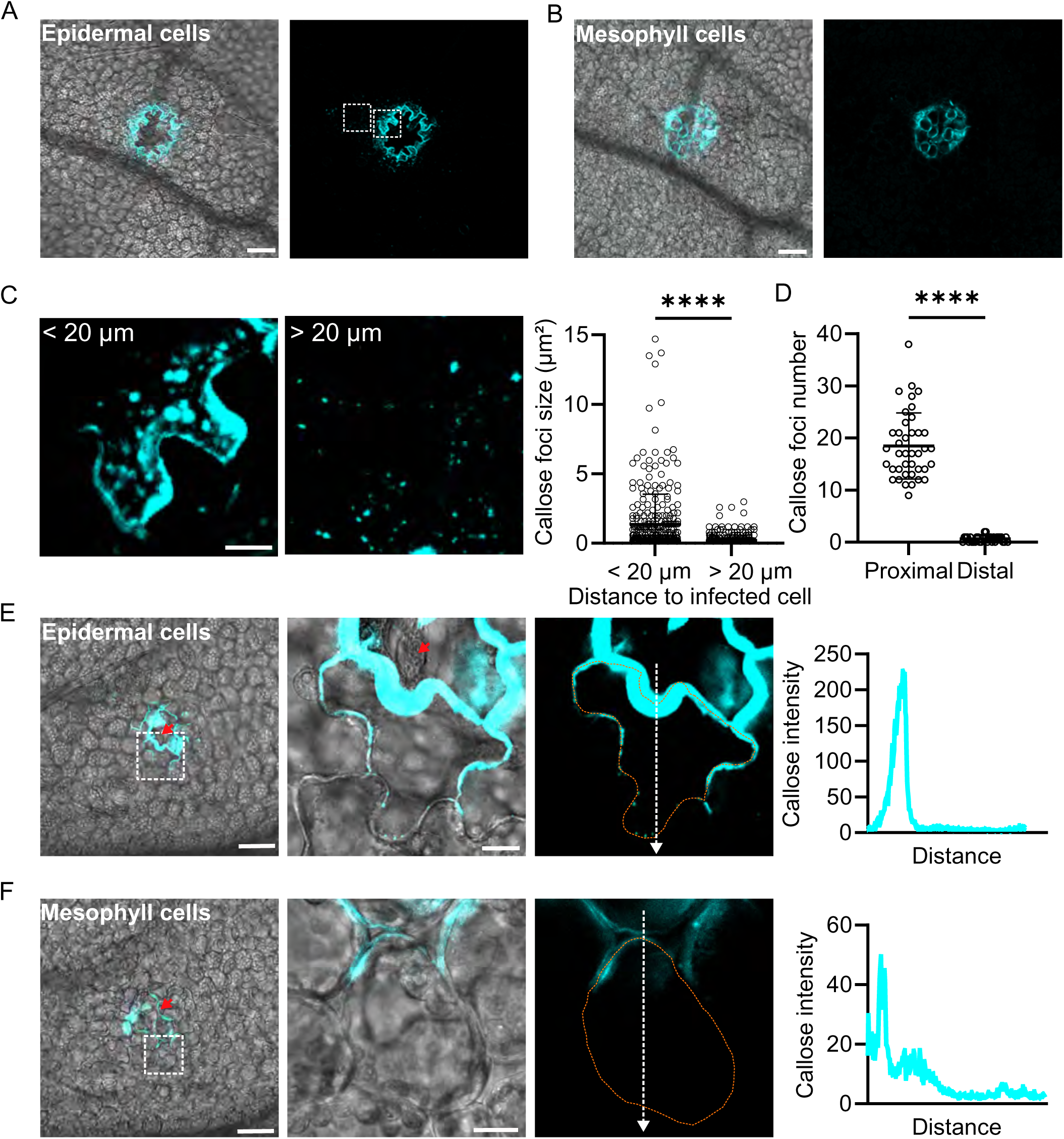
Spatially restricted deposition of callose at the infection front after ETI activation. Two-week-old *Arabidopsis* seedlings were flood-inoculated with *Pst* DC3000 (AvrRpm1) or mock treated with 5 mM MgCl₂. Callose deposition was visualized by aniline blue staining and imaged by confocal microscopy. **(A-B)** Callose deposition in cells proximal to infection sites. Both epidermal (A) and mesophyll (B) cells exhibit localized callose accumulation at 48 hpi. Mesophyll cells in (B) are directly underneath of the epidermal cells in (A). Scale bar, 50 μm. Insets are shown in (C). **(C)** Size (µm²) of callose foci in proximal (<20 µm) and distal (>20 µm) regions of infection sites. Representative insets are shown on the left, and quantification is shown on the right. Scale bar, 10 µm. **(D)** Quantification of callose foci per 400 µm² in proximal and distal regions relative to infection sites. Proximal regions are defined as cells within five cell layers of the infection site, whereas distal regions are defined as cells located more than five cell layers away. **(E-F)** Callose deposition displays a polarized distribution in cells adjacent to infection sites, with higher accumulation on the side facing the infection and lower accumulation on the opposite side. This polarized pattern is observed in both epidermal (E) and mesophyll (F) cells. Infection sites are determined by bacterial colonization near stomata, as indicated by red arrows. Scale bar, 50 µm and inset bars, 10 µm. Mesophyll cells in (F) are right underneath the epidermal cells in (E). Line-scan analyses extending outward from the infiltration site quantify the spatial distribution of callose intensity.

Beyond its spatial restriction, callose deposition within cells adjacent to bacterial colonies exhibited intracellular polarization. In both epidermal and mesophyll cells, callose accumulated asymmetrically, with higher accumulation on the cell wall face oriented toward the infection site and markedly lower accumulation on the distal face (Fig. 3E-F). Line-scan analyses confirmed this polarized distribution, with signal intensity peaking at the pathogen-proximal wall and declining sharply toward the opposite side of the cell. This polarization was observed in both epidermal cells and in underlying mesophyll cells, indicating that the polarity signal propagates across cell layers. Together, these results show that callose deposition is confined to a defined population of cells at infection sites and displays both tissue-level and subcellular spatial organization during plant immune responses.

### SA and JA biosynthesis genes are expressed in spatially distinct, adjacent cell populations

Plants activate multiple defense hormone pathways upon pathogen recognition, including the antagonistic SA and JA signaling networks (26, 43–45). Although SA–JA antagonism has been extensively characterized at the molecular level, whether this antagonism operates within individual cells or emerges through spatial organization across neighboring cell populations has remained unclear. To address this question, we generated a dual fluorescent reporter line co-expressing transcriptional fusions to the promoters of *CBP60g* (Cerulean), a key transcriptional regulator of SA biosynthesis, and *ALLENE OXIDE SYNTHASE* (*AOS*, mVenus), a core enzyme in JA biosynthesis, enabling simultaneous visualization of SA- and JA-associated transcriptional responses at single-cell resolution (14, 24, 38, 39, 46–48).

We first validated the dual reporter line by comparing its behavior to previously characterized single-reporter lines. *CBP60g* expression was spatially restricted around infection sites and sustained at later time points, consistent with our previously published *CBP60g* single-reporter line infected with *Pst* DC3000 (AvrRpm1) (Fig. 4A, S3A-B) (14). *AOS* expression was similarly induced following infection and persisted over time, consistent with results from a previously published *AOS* single-reporter line (Fig. 4A–B, Fig. S3C) (46). Notably, *AOS* expression bundles were broader than *CBP60g* bundles and frequently merged at later infection stages, making discrete boundaries less distinguishable (Fig. 4A).

**Figure 4.**
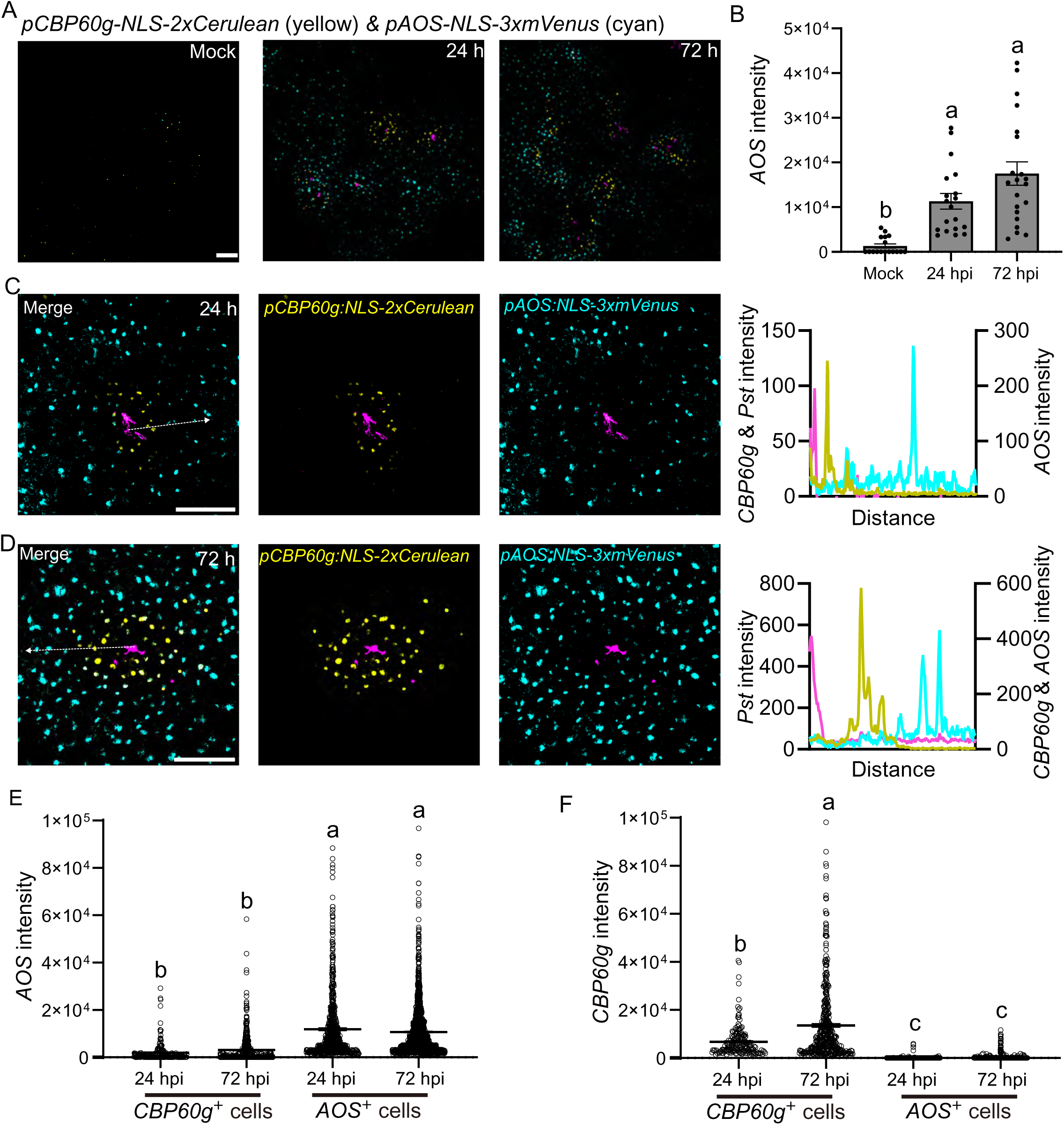
Spatial separation of SA and JA biosynthesis gene expression at bacterial infection sites. Two-week-old *Arabidopsis* seedlings expressing dual reporters *pCBP60g::NLS–2xCerulean* (SA biosynthesis) and *pAOS::NLS–3xmVenus* (JA biosynthesis) were flood-inoculated with *Pst* DC3000 (AvrRpm1) or mock treated with 5 mM MgCl₂. *CBP60g* and *AOS* expression are shown in yellow and cyan, respectively. Bacteria are shown in pink. Images are maximum projections of confocal z-stacks. **(A-B)** Expression of *CBP60g* and *AOS* at bacterial infection sites. Left panels show reporter expression at infection sites. Scatter plots show quantification of mVenus fluorescence intensity per image. Total fluorescence intensity was measured, and scatter plots show the median ± SD (n ≥ 6 images from at least 3 plants). Different letters indicate statistically significant differences (*p* < 0.0001, ANOVA with Tukey test). Scale bar, 100 μm. **(C-D)** Spatial relationship between *CBP60g* and *AOS* expression domains at infection sites. *AOS* expression is mainly suppressed in *CBP60g*-expressing cells but strongly induced in adjacent domains outside the *CBP60g* expression region. **(E-F)** Quantification of *CBP60g* and *AOS* expression in *CBP60g*- or *AOS-*expressing cells (*CBP60g⁺* and *AOS⁺* cells) at 24 and 72 hpi. *CBP60g* and *AOS* expression are primarily detected in distinct cells. *CBP60g* reporter intensity increases at 72 hpi. Total fluorescence intensity was measured, and scatter plots show the median ± SD (n ≥ 6 images from at least 3 plants). Different letters indicate statistically significant differences (*p* < 0.0001, ANOVA with Tukey test).

Next, we examined whether SA- and JA-associated transcriptional responses occur within the same cells or are spatially segregated during ETI-potentiated immunity. High-magnification imaging at infection sites with clearly defined bacterial colonies revealed two recurring expression patterns at 24 hpi (Fig. 4C, S4A). In the first pattern, *CBP60g* and *AOS* were both induced in cells immediately adjacent to bacterial colonies, yet their expression was largely mutually exclusive at the single-cell level: cells with high *CBP60g* expression exhibited low *AOS* signal, whereas immediately adjacent cells displayed the opposite pattern (Fig. S4A). This side-by-side arrangement indicates that SA- and JA-associated activities are partitioned between neighboring cells even at early infection stages. In the second pattern, which was most frequently observed at 24 hpi, *CBP60g* and *AOS* expression formed clearly delineated spatial domains at infection sites (Fig. 4A-C). *CBP60g* expression was highest in cells surrounding bacterial colonies, defining a spatially restricted inner domain, while *AOS* expression was elevated in cells located outside this region, forming a surrounding domain. Cells at the interface between these domains showed relatively low expression of both reporters. By 72 hpi, this SA and JA domain-based organization became more prominent, with most infection sites exhibiting clearly separated *CBP60g*- and *AOS*-enriched regions (Fig. 4D). Quantitative analysis confirmed that *AOS* expression was significantly lower in *CBP60g*⁺ cells than in *AOS*⁺ cells, and *CBP60g* expression was correspondingly reduced in *AOS*⁺ cells, consistent with mutual exclusivity of the two biosynthetic programs across neighboring cell populations (Fig. 4E–F).

### Spatially compartmentalized SA–JA antagonism shapes tissue-level defense architecture

Then, we asked whether downstream hormone response outputs exhibit distinct spatial behaviors. To test whether the spatial arrangement observed for SA and JA biosynthesis reporters is propagated to downstream response genes, we examined expression of the canonical SA response marker *PR1* and the JA response marker *Vegetative Storage Protein 1* (*VSP1*) (29, 49–51). Using the previously published *pPR1::NLS-3xVenus* reporter line (46, 52), we generated *PR1*-*VSP1* dual reporter plants to enable simultaneous visualization of SA and JA response domains. We first validated the specificity of this dual reporter system. SA spray treatment strongly induced *PR1* but not *VSP1* expression, whereas mechanical damage induced *VSP1* but not *PR1* (Fig. S5). These results confirm the specificity of the dual reporter line.

Using this system, we analyzed the expression of *PR1* and *VSP1* in four-week-old *Arabidopsis* following infiltration with a high inoculum (OD600 = 0.2) of *Pst* DC3000 (AvrRpm1). At both 24 and 72 hpi, *PR1* and *VSP1* retained a similar overall spatial organization, with an inner *PR1* domain adjacent to the infection site and an outer *VSP1* domain (Fig. 5A–B). Line-scan analyses extending outward from the infiltration site revealed suppressed expression of both markers at the infiltrated regions, followed by mutually exclusive expression of *PR1* and *VSP1* in surrounding regions (Fig. 5C-D). To systematically compare *PR1* and *VSP1* expression across space and time, we defined three response regions relative to the infiltration border: proximity I (P I, <0.6 mm), proximity II (P II, 0.6-1.2 mm), and distal (D) regions, based on previously reported domain boundaries (Fig. 4E) (29). At 24 hpi, *PR1* expression peaked in P I, the region immediately adjacent to bacterial infection. By 72 hpi, *PR1* expression remained high in P I but declined significantly in P II and distal regions (Fig. 5F). In contrast, *VSP1* expression was lowest in P I and highest in P II at 24 hpi, reinforcing the spatial exclusivity between SA and JA response outputs (Fig. 5G). At later stages, *VSP1* expression decreased significantly in both P I and distal regions, indicating dynamic remodeling of JA responses over time. In very distal leaf tip regions, *VSP1* expression was high while *PR1* expression remained low within the same cells, indicating the establishment of local acquired resistance throughout the leaf (Fig. 5H). Collectively, these findings reveal that SA-JA responses are spatially compartmentalized and dynamically regulated across the leaf, with SA-related activities enriched near the infection front and JA-related activities predominating in surrounding regions. This organization provides a spatial framework for coordinating multiple defense programs.

**Figure 5.**
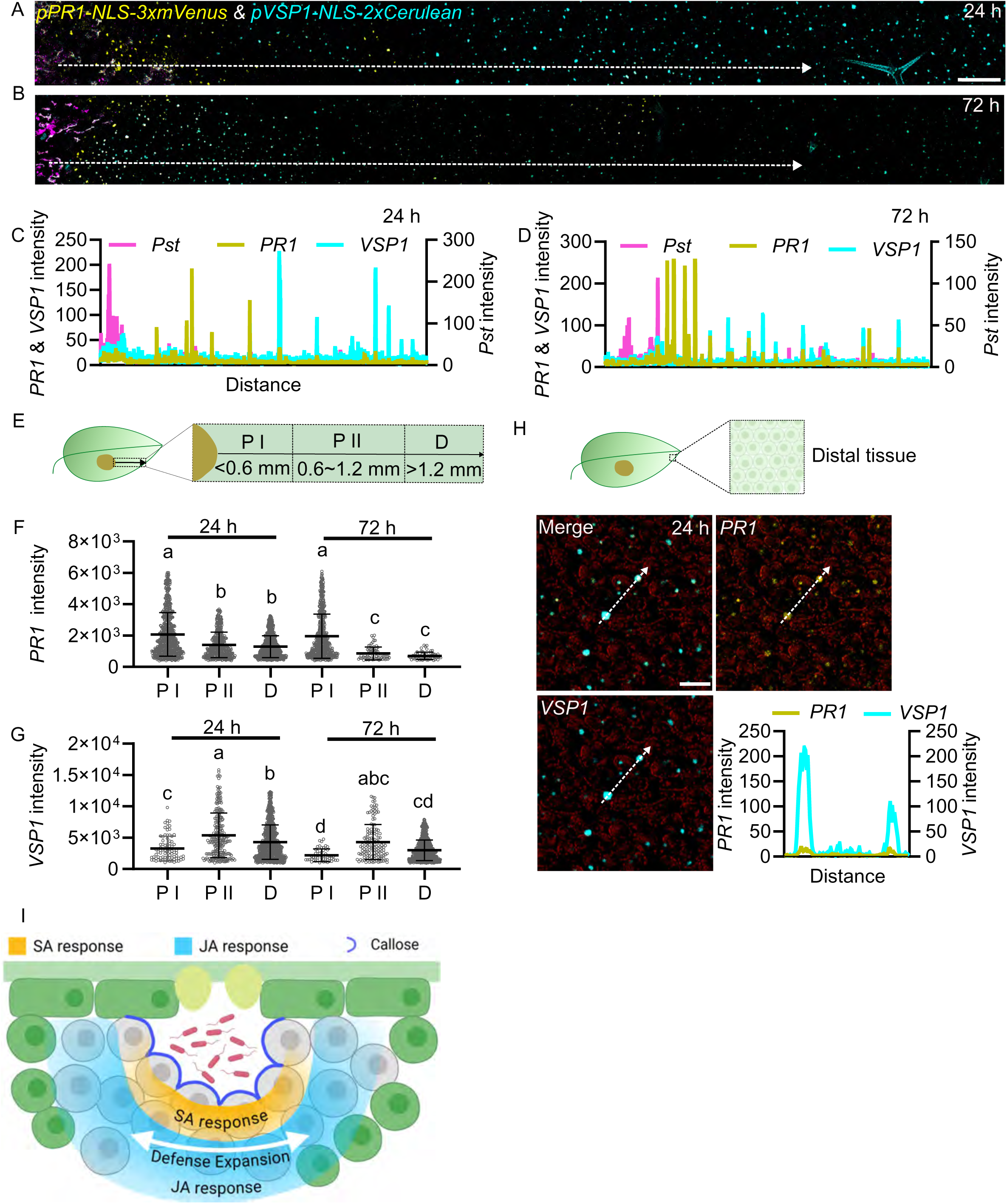
Spatially organized SA-JA antagonism shapes defense responses during bacterial infection. Four-week-old *Arabidopsis* leaves expressing dual reporters *pPR1::NLS–3xmVenus* (SA response) and *pVSP1::NLS–2xCerulean* (JA response) were infiltrated with *Pst* DC3000 (AvrRpm1) at a suspension of 1× 10^8^ CFU/mL. *PR1* and *VSP1* expression are shown in yellow and cyan, respectively. Bacteria are shown in pink. Images are maximum projections of confocal z-stacks. **(A-B)** Spatial organization of *PR1* and *VSP1* expression at infection sites at 24 and 72 hpi. *VSP1* expression is suppressed in *PR1*-expressing cells but strongly induced in adjacent domains outside the *PR1* region. Scale bar, 0.5 mm. **(C-D)** Line-scan analysis across the regions indicated in (A-B) showing the spatial distribution of bacteria, *PR1*, and *VSP1* fluorescence signals. **(E)** Schematic diagram of defined response regions relative to the infiltration border: proximity I (P I), proximity II (P II), and distal (D). The infiltrated area is shown in brown. **(F-G)** Quantification of *PR1* and *VSP1* fluorescence intensity in defined response regions shown in (E) at 24 and 72 hpi. Total fluorescence intensity density per region was measured. Scatter plots show median ± SD (n = 6 regions from 3 images of 3 plants). Different letters indicate statistically significant differences (*p* < 0.0001, ANOVA with Tukey test). Experiments were repeated twice with similar results. **(H)** Representative images of distal tissue regions showing high *VSP1* expression and low *PR1* expression within the same cells. Scale bar, 50 μm. Line-scan analysis shows fluorescence intensity profiles of *PR1* and *VSP1* across the indicated region. **(I)** Model illustrating spatially organized plant defense at the infection front.

## Discussion

Effector-triggered immunity (ETI) has been historically viewed as a strong immune response associated with cell death and pathogen killing. Increasing evidence indicates that ETI outputs are spatially constrained and tightly regulated rather than uniformly deployed across infected organs (13, 15, 53). Here, we show that ETI organizes plant defense into a spatially structured containment architecture (Fig. 5I). Using live-cell reporters together with fluorescently labeled bacteria, we demonstrate that immune activation persists for days yet remains confined to a narrow zone of cells surrounding viable bacterial colonies (Fig. 1). Within this zone, defense responses are dynamically reorganized over time, with transcriptional activation initially peaking in cells immediately adjacent to bacteria and subsequently shifting outward into neighboring cell layers (Fig. 2). Together with localized callose deposition and hormone-specific transcriptional domains, these findings reveal a multilayered immune boundary that contains pathogen spread (Fig. 5I).

Spatial restriction is increasingly recognized as a feature of plant immune responses. In roots, immune activation can be confined to specific cell layers or regions following MAMP perception or damage-associated signals (9, 18, 54). Local immune responses also trigger spatially restricted signaling gradients, including Ca²⁺, ROS, iron availability, and SA accumulation (28, 46, 55, 56). ETI-potentiated immunity also induces prolonged activation of SA-related defense markers that are transiently observed during PTI (14). Recent spatial and single-cell analyses have identified the presence of specialized immune-active and immune-supporting cell populations, including rare primer cells and neighboring bystander cells (15, 16). Our results extend these findings by linking cellular responses to precise pathogen position, revealing that leaf immune responses are organized into discrete, pathogen-centered immune domains.

In addition to spatial restriction, immune signaling spreads between neighboring cells to reinforce defense at infection fronts. Using *FRK1* as a marker of immune activation, we observed strong and sustained induction that remained limited to a few cell layers surrounding each bacterial colony (Fig. 1 and S1). Notably, the peak of *FRK1* expression shifted outward over time, indicating propagation of immune activation to adjacent cells (Fig. 2). Cells directly contacting bacteria undergo cell death as a result of ETI, whereas surrounding cells likely receive propagated defense signals such as ROS, Ca²⁺, DAMPs, phytocytokines, and apoplastic pH changes that reinforce immunity (13, 57, 58). Similar short-range propagation of immune signaling has been observed in roots and leaves (18, 46, 56), suggesting that intercellular signaling networks coordinate defense activation across neighboring cells. This immune propagation is consistent with the model of localized acquired resistance (LAR). The LAR model can now be refined to highlight immune activation is restricted to a localized ring of cells surrounding infection sites within a leaf, forming a defensive boundary that prevents systemic spread (13, 59–61).

Our data further highlight that immune activation limits pathogen spread but does not kill bacteria already present at infection sites. ETI induced localized defense responses, including defense gene activation, callose deposition, and hormone signaling, that reinforced defenses around bacterial microcolonies. Such localized reinforcement likely limits bacterial spread by strengthening physical barriers, such as oxidative cross-linking, callose deposition, and lignin deposition (17, 62–65). Despite this strong physical and cellular response, bacterial populations remained viable within these confined regions, as determined by bacterial fluorescence and colony counts of total bacterial populations in leaf tissue. These observations are consistent with previous studies of bacterial and viral infections, where immune activation reduces pathogen spread but high titers of infectious pathogens can be recovered from tissue undergoing programmed cell death (13, 66–69). Together, these findings support a model in which ETI functions primarily as a spatial containment strategy, restricting pathogen expansion and preventing the establishment of new infection foci rather than achieving immediate pathogen eradication.

The establishment of spatially confined immune domains is accompanied by coordinated phytohormone patterning. SA and JA signaling pathways are classically described as antagonistic, raising the question of how they are coordinated during immune responses (70). Using dual reporter lines, we observed a clear spatial separation between SA- and JA-associated responses, with SA markers enriched in cells adjacent to bacterial colonies and JA markers predominantly expressed in surrounding cells (Fig. 4–5). Similar spatial segregation of SA and JA activity has been reported around infection sites single-reporter studies and recent SA biosensor analyses (28–30), suggesting that antagonistic pathways can be deployed in parallel through tissue-level organization. This organization supports a division of labor in which distinct cell populations execute complementary defense functions in a coordinated defense domain.

In summary, our findings support a model in which LAR operates through spatially organized defense domains that form around infection sites. Within these domains, immune activation propagates and hormone responses are partitioned to coordinate distinct cellular responses. This organization highlights ETI as a localized containment strategy rather than a uniformly deployed response. Understanding how these domains are established and integrated with systemic immunity will be an important direction for future work.

## Materials and Methods

### Plant materials and growth conditions

*Arabidopsis thaliana* ecotype Columbia (Col-0) and derived fluorescent reporter lines were used in this study. The transcriptional reporter lines *pFRK1::NLS-3xmVENUS*, *pPR1::NLS-3xmVENUS*, and *pAOS::NLS-3xmVENUS* were kindly provided by Professor Niko Geldner (18, 46). Seeds (Col-0 or transgenic lines) were surface-sterilized with 50% bleach containing 0.1% Tween 20 for 8 min, followed by 75% ethanol for 1 min, and rinsed four times with sterile water before plating on half-strength (½) Murashige and Skoog (MS) medium. Seeds were stratified in the dark at 4 °C for 2 days prior to germination on soil or ½ MS medium. Ten- to fourteen-day-old seedlings grown on ½ MS medium were maintained at 22 °C under long-day conditions (16 h light/8 h dark) and used for microscopy analyses. For pathogen infection assays and plant transformation, four-week-old plants grown in soil were maintained in a controlled environment chamber at 22 °C and 70% relative humidity under a 10 h light/14 h dark photoperiod (100 μmol m⁻² s⁻¹).

### Bacterial strains and growth conditions

*Pseudomonas syringae* pv. tomato DC3000 (*Pst* DC3000) was labeled with 3xmCherry (*att*Tn*7*-3xmCherry) using a site-specific Tn7 3xmCherry vector to generate *Pst* DC3000-mCherry (14). *Pst* DC3000-mCherry expressing AvrRpm1, AvrRpt2, or AvrRps4 were generated using broad host range plasmid pVSP61, respectively (71). Transformants were selected on nutrient yeast glycerol agar (NYGA) medium supplemented with 100 μg/mL rifampicin, 50 μg/mL spectinomycin, and 50 μg/mL kanamycin, and colonies were screened after 2 days of incubation at 28 °C. For all inoculation experiments, bacterial strains were grown overnight at 28 °C on NYGA medium containing the same antibiotic combination prior to preparation of inoculum.

### Bacterial inoculation and quantification

For inoculation assays, *Pst* DC3000-3xmCherry from overnight cultures was harvested and resuspended in 5 mM MgCl₂ to the indicated optical density (OD600). For seedling flood inoculation, two-week-old plants grown on ½ MS medium were submerged in 40 mL of bacterial suspension (OD600= 0.0005) supplemented with 0.02% Silwet L-77 per 100 × 100 mm square Petri dish (Fisherbrand). After 20-30 s at room temperature, excess bacterial suspension was removed, and plates were sealed with 3M Micropore tape before incubation in a growth chamber. For mature plant infections, fully expanded leaves of four-week-old *A. thaliana* were inoculated either by syringe infiltration (OD600 = 0.0002 or 0.2) or by spray inoculation (OD600 = 0.2). Following inoculation, plants were maintained under ambient humidity for 1 h to allow evaporation of excess surface moisture, then covered with a transparent dome to maintain high humidity and incubated in a growth chamber for 24 - 72 h.

Bacterial growth following flood inoculation was quantified as colony-forming units (CFU) per milligram of tissue. For each biological replicate, roots were removed from three seedlings and the aerial tissues were pooled (5-7 replicates per time point). After recording fresh weight, samples were homogenized in 5 mM MgCl₂, serially diluted, and plated on NYGA medium supplemented with 100 μg/mL rifampicin, 50 μg/mL spectinomycin, and 50 μg/mL kanamycin. Colonies were counted after incubation at 28 °C.

### Generation of transgenic lines

Approximately 2 kb promoter regions upstream of the start codon of *CBP60g* (AT5G26920; 2183 bp) and *VSP1* (AT5G24780; 2333 bp) were PCR-amplified from genomic DNA of *A. thaliana*. Promoter fragments were cloned into pENTR vectors and fused to a nuclear localization signal (NLS) using the In-Fusion HD Cloning Plus system (Clontech). Primers for *CBP60g* and *VSP1* are F*_CBP60g_*-CGACTCTAGAGGATCTGGCTCGATCAAACTTAGATATCAA, R*_CBP60g_*-TGGCATCCATGGATCCATGATCACTTTTAGGTTTAGAGAA and F*_VSP1_*-CGACTCTAGAGGATCTATGACCCACGACCAGAT, R*_VSP1_*-TGGCATCCATGGATCCATTTTTTTGTATGGTTTATTGTTT. Entry clones were subsequently recombined into the binary destination vector pMpGWB123 containing 2xmCerulean via Gateway cloning (Invitrogen) (72). Constructs were introduced into *Agrobacterium tumefaciens* strain GV3101 and transformed into *A. thaliana* p*AOS*- and p*PR1*-fluorescent reporter lines using the floral dip method. Transgenic seedlings were selected on ½ MS medium supplemented with 50 mg/L kanamycin and 100 mg/L carbenicillin. Ten to fifteen independent T1 lines were screened for mCerulean fluorescence, and three to five T2 lines exhibiting stable expression were selected for bacterial infection assays. T2 or T3 plants (1–2 independent lines) displaying consistent induction patterns following bacterial infection were used for subsequent experiments.

### Callose staining

Callose deposition in *Arabidopsis* leaves was visualized using aniline blue staining (73, 74). Whole seedlings were harvested at 48 h post-inoculation and cleared in ethanol: acetic acid = 3:1 (v:v) for 1 h at 37⁰C until chlorophyll was completely removed. Samples were subsequently rehydrated with 70%, 50%, and 30% ethanol series for 30 min at each step. Cleared seedlings were rinsed once in 150 mM K₂HPO₄ buffer (pH 9.5). Samples were then stained with 0.01% (w/v) aniline blue prepared in 150 mM K_2_HPO_4_ buffer (pH 9.5) for 1 h in the dark. Stained tissues were mounted in 50% glycerol and imaged using a confocal microscope.

### Confocal settings and image processing

Confocal imaging was performed using either a Leica TCS SP8 or a Leica TCS SP8 STED/MP confocal microscope (Leica Microsystems). Images were acquired with 10x and 20x dry objectives, and tile scans were collected using a 5x objective with 10% overlap. The following excitation and emission settings were used on the Leica TCS SP8: mVENUS/mCitrine (excitation 488 nm; emission 493–540 nm), mCherry (552 nm; 586–635 nm), mCerulean (405 nm; 433–496 nm), chlorophyll autofluorescence (638 nm; 650–720 nm), and callose (405 nm; 433–480 nm); and on the Leica TCS SP8 STED/MP system: mCitrine (488 nm, 500–551 nm), mCherry (587 nm, 599–649 nm), mCerulean (458 nm, 466–501 nm), chlorophyll (488 nm, 650–725 nm). Sequential scanning was applied to minimize channel crosstalk. For time-course imaging of transcriptional reporter lines, all images within each experiment were acquired under identical settings, including objective lens, laser power, pinhole size, detector gain, and Z-stack intervals, to ensure accurate comparison of fluorescence intensity over time. Imaging parameters were optimized for each reporter line based on transgene expression levels but remained consistent between treatments within the same experiment.

Fluorescence intensity, colony size, and callose deposition were quantified using the Fiji image processing package (http://fiji.sc/Fiji). All images within each experiment were processed using identical parameters. For fluorescence quantification, z-stack images were first projected by summing slices for each channel. A consistent gray value threshold was then applied within each experiment to distinguish true signal from background; for example, signals below a gray value of 150 were excluded to minimize autofluorescence and weak background signals. Threshold values were optimized for each reporter line based on signal intensity but remained constant between mock and treatment groups within the same experiment. To reduce noise, a minimum particle size threshold was defined before quantifying the total signal area. Total fluorescence intensity for each nucleus or defined region of interest (ROI) was calculated as the product of the measured signal area and the mean pixel intensity within that area.

Line-scan analyses were performed using Leica Application Suite X (LAS X, Leica Microsystems). Linear ROIs were drawn across selected cells or tissue regions to generate fluorescence intensity profiles along the defined axis. Background subtraction, when required, was applied uniformly across samples, and identical acquisition and analysis settings were maintained to ensure comparability between treatments.

### Quantification and Statistical Significance

Statistical analyses were conducted using GraphPad Prism 10.0 (https://www.graphpad.com/). Data are presented as mean ± standard deviation (SD), with *n* denoting the number of analyzed images collected from at least three independent plants. One-way analysis of variance (ANOVA) followed by Tukey’s post hoc test was used for multiple comparisons, while two-tailed Student’s *t*-tests were applied for pairwise comparisons. Additional details regarding statistical analyses are provided in the figure legends.

## Data, Materials, and Software Availability (data sharing plans (for all data, documentation, and code used in analysis))

Raw data in the form of microscopy images have been deposited in Zenodo (https://doi.org/10.5281/zenodo.19358730).

Seeds and plasmids will be deposited in Arabidopsis Biological Resource Center (ABRC) and Addgene, respectively, and made available upon publication.

## Acknowledgments and funding sources

We thank Dr. Niko Geldner for generously providing *FRK1, PR1, and AOS* transcriptional reporter seeds. We are grateful to Dr. Pamela Ronald and the Advanced Light Microscopy Core at the University of Missouri for access to confocal microscopy facilities and technical support. We also thank members of the Coaker lab and Zhu lab for valuable discussions and input throughout the course of this work. This project was supported by a grant from the NIH (R35GM136402) to G.C., as well as the startup grant and Research Council Grant to J.Z. from the University of Missouri.

## Author contributions

J.Z. and G.C. conceived and conceptualized the project; X.L, Y.S. and J.Z. designed, performed experiments and analyzed data; J.Z. and G.C. wrote the original draft; X.L, Y.S., G.C. and J.Z reviewed and edited the manuscript. G.C. and J.Z supervised the project and acquired funding.

