## Supplemental figures for "Spatial organization of plant defense at the infection front"

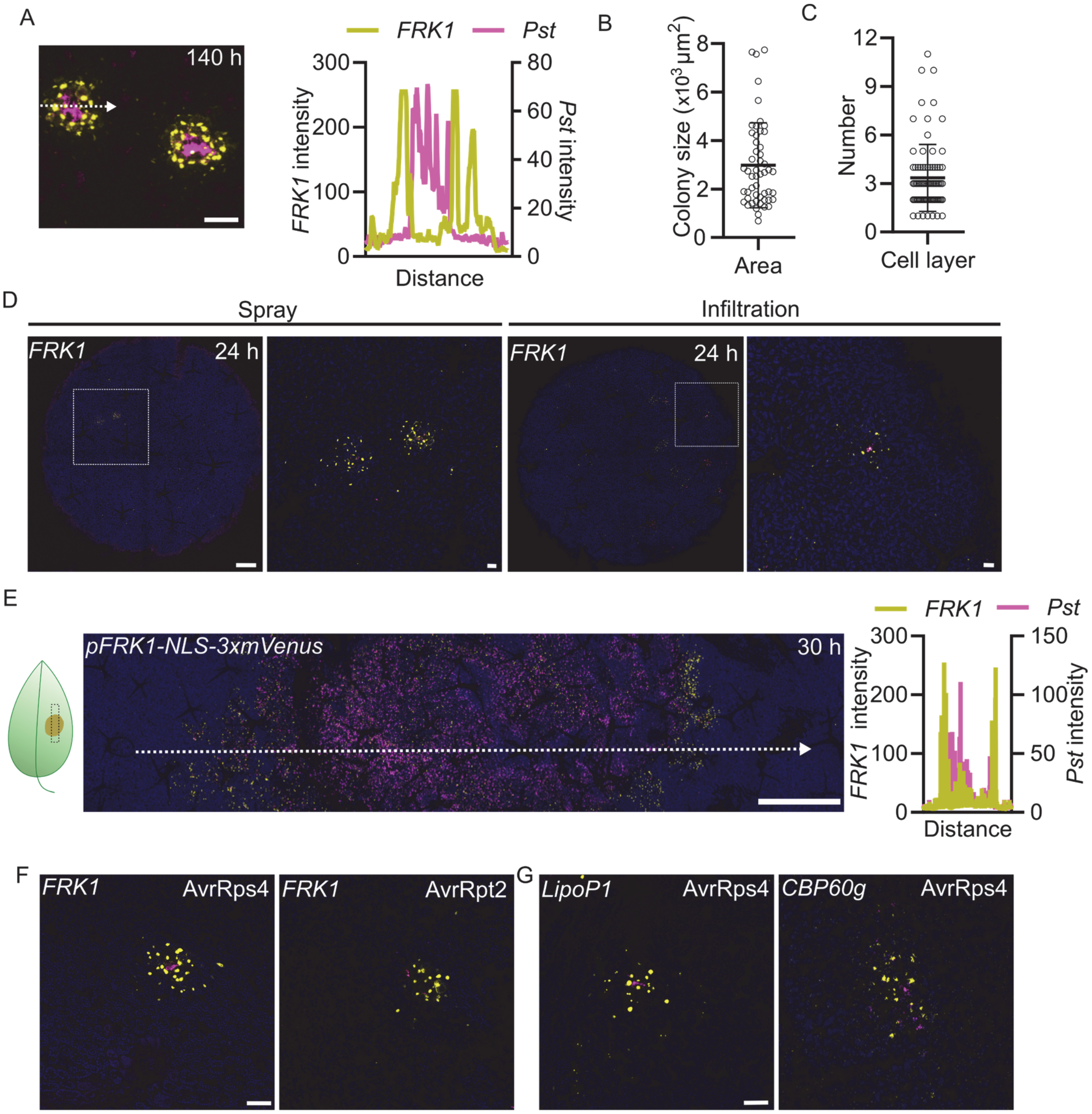


**Fig. S1.** Sustained and spatially restricted *FRK1* expression in *Arabidopsis* leaves

(A) Two-week-old *Arabidopsis* seedlings expressing *pFRK1::NLS–3xmVenus* were flood-inoculated with *Pst* DC3000 (AvrRpm1) expressing mCherry or mock treated with 5 mM $MgCl_2$. Images are maximum projections of confocal z-stacks. Chlorophyll autofluorescence is shown in cyan. Representative confocal image showing spatially restricted *FRK1* expression at 140 hpi following ETI activation. Image is a maximum projection of confocal z-stacks. Scale bar, 100 μm.

(B) Quantification of bacterial colony size at 140 hpi.

(C) Quantification of the number of *FRK1*-activated cell layers at 140 hpi.

(D) Spatially restricted *FRK1* expression following both spray inoculation and syringe infiltration on mature *Arabidopsis*. Leaves of four-week-old *Arabidopsis* were sprayed with *Pst* DC3000

(AvrRpm1) at 1x $10^8$ colony-forming units/mL (CFU/mL, OD600=0.2) or infiltrated with OD600=0.0002. Representative images are maximum projection from confocal z stacks. Scale bar, 200 μm. Inset bar, 50 μm.

(E) High-density infiltration of avirulent bacteria induces spatially restricted *FRK1* responses in mature leaves. Four-week-old *Arabidopsis* plants were infiltrated with *Pst* DC3000 (AvrRpm1) at OD600 = 0.2. Left: representative confocal image (maximum projection of z-stacks). Scale bar, 0.5 mm. Right: quantification of *FRK1* response and bacterial growth.

(F) Spatial restriction of *FRK1* expression after both AvrRpt2 and AvrRps4-triggered immunity at 48 h after flood inoculation of seedlings. Scale bar, 50 μm

(G) Expression of immune related genes *CBP60g* and *LipoP1* shows similar spatial restriction pattern at 48 h after flood inoculation of seedlings. Scale bar, 50 μm.

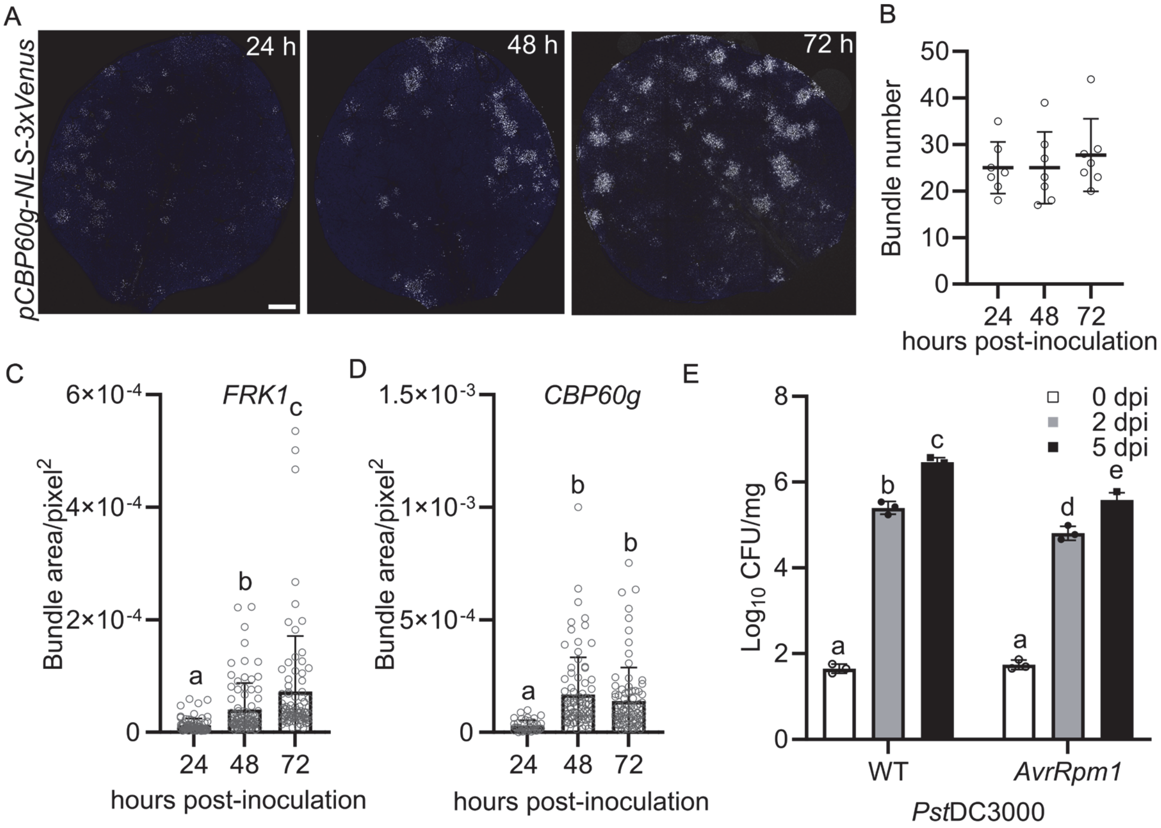


**Fig. S2.** ETI activation suppresses subsequent bacterial invasion.

Two-week-old *Arabidopsis* seedlings expressing p*FRK1*::NLS–3xmVenus or p*CBP60g*::NLS–2xCerulean were flood-inoculated with Pseudomonas syringae pv. tomato (*Pst*) DC3000 (AvrRpm1) expressing mCherry or mock treated with 5 mM $MgCl_2$. Images are maximum projections of confocal z-stacks.

(A) Representative images of *CBP60g* expression bundles over time following ETI activation. Images are maximum projections of confocal z-stacks. *CBP60g* signal is shown in white and chlorophyll autofluorescence in blue. Scale bar, 0.5 mm.

(B) Quantification of *CBP60g* expression bundle numbers shown in (A). Six images from at least three plants were analyzed. No statistically significant differences were detected (ANOVA with Tukey test).

(C–D) Quantification of *FRK1* and *CBP60g* bundle areas over time. Six images from at least three plants were analyzed. Different letters indicate statistically significant differences ($p < 0.0001$, ANOVA with Tukey test). Experiments were repeated twice with similar results.

(E) Growth of *Pst* DC3000 and *Pst* DC3000 (AvrRpm1) in *Arabidopsis* (Col-0) seedlings flood-inoculated with $2.5 \times 10^5$ CFU/mL (OD600 = 0.0005) bacterial suspension.

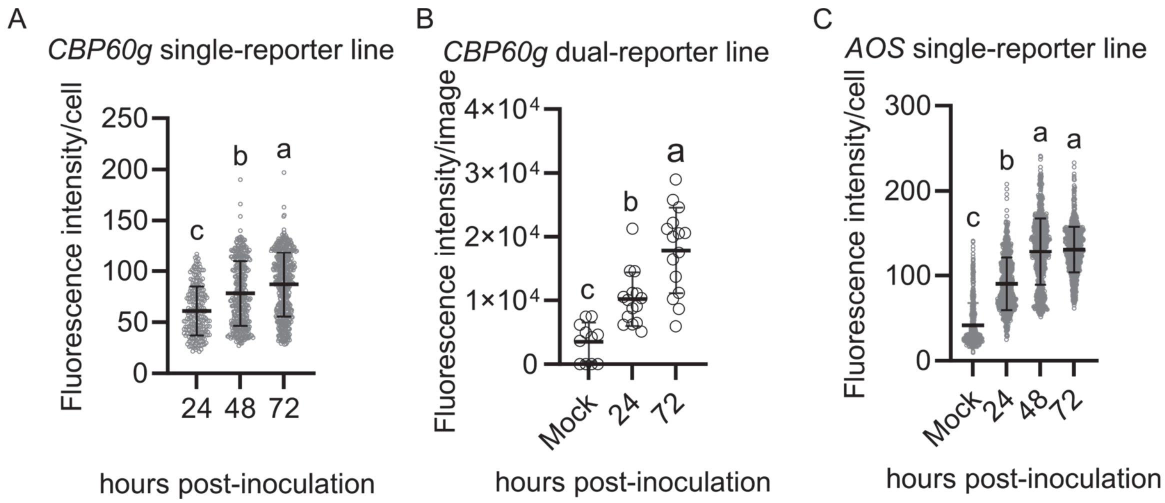


**Fig. S3.** Consistent *CBP60g* and *AOS* expression patterns in both single- and double-reporter lines.

Two-week-old *Arabidopsis* seedlings grown on MS plates were flood-inoculated with with *Pst* DC3000 (AvrRpm1) expressing mCherry (OD600 = 0.0005) or mock treated with 5 mM $MgCl_2$. (A–C) Quantification of fluorescence intensity for *CBP60g* and *AOS* reporters. Total fluorescence intensity density per nucleus (A, C) and per image (B) was measured. Scatter plots show median ± SD (n ≥ 6 images from at least 3 plants). Different letters indicate statistically significant differences ($p$ < 0.0001, ANOVA with Tukey test).

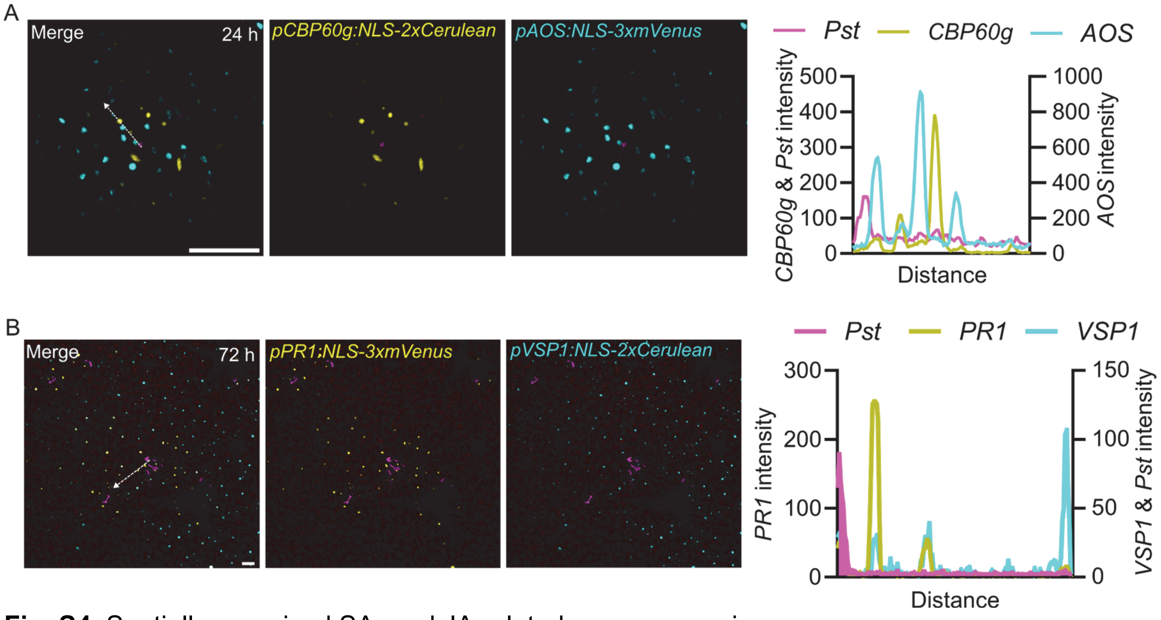


**Fig. S4.** Spatially organized SA- and JA-related gene expression.

(A) Spatially separated induction of *AOS* and *CBP60g* near bacterial infection sites at early stages. Left: representative confocal images (maximum projections). Right: line-scan quantification corresponding to the dashed line in the merged image. Distance represents increasing distance from the bacterial colony. Scale bar, 100 μm.

(B) Spatial organization of *PR1* and *VSP1* responses in *Arabidopsis* seedlings following flood inoculation with *Pst* DC3000 (AvrRpm1). Two-week-old seedlings were inoculated with bacterial suspension at OD600 = 0.0005.

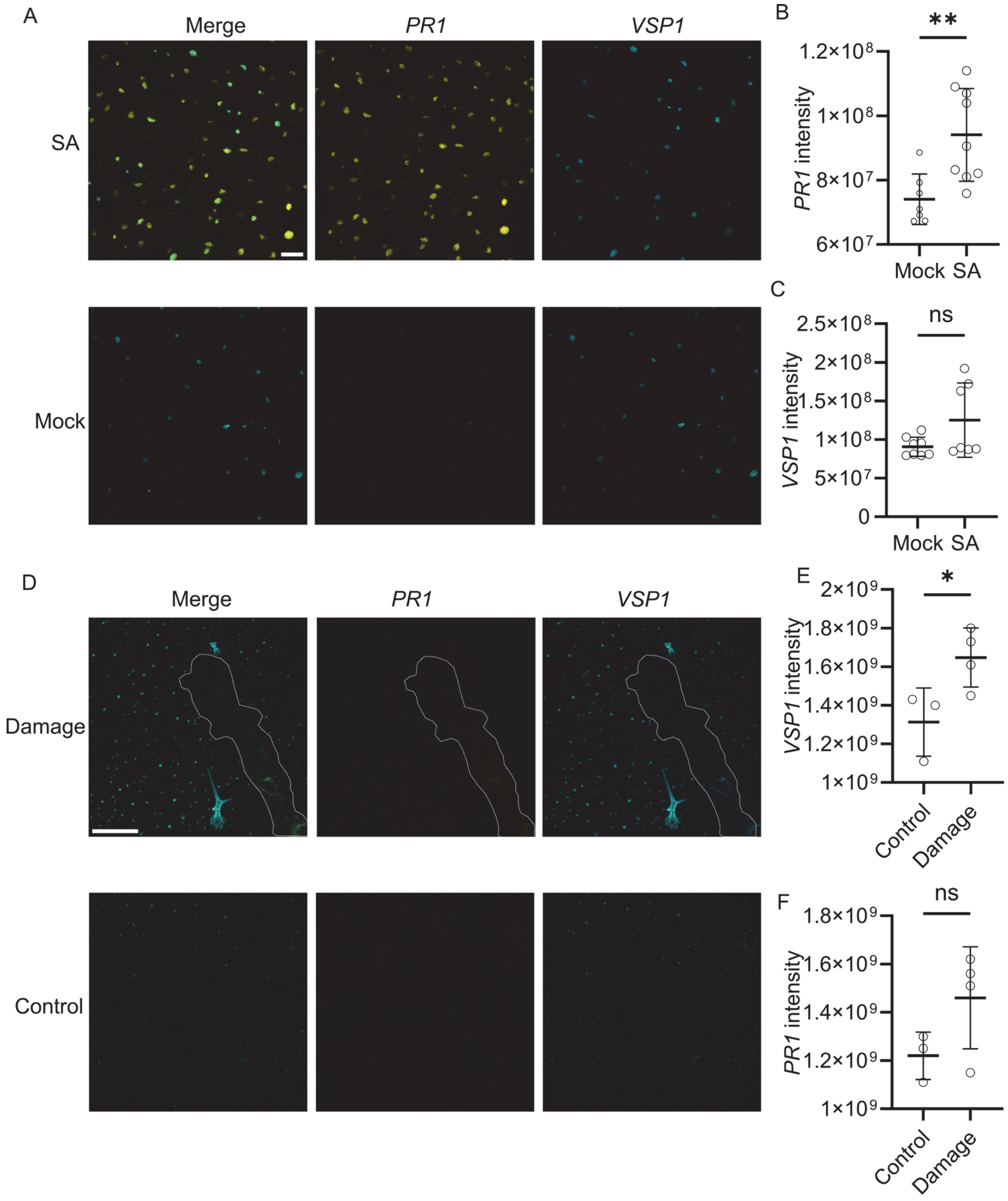


**Fig. S5.** Validation of the *PR1*-*VSP1* dual-reporter line.

Leaves of four-week-old *Arabidopsis* plants expressing *PR1* and *VSP1* reporters were treated with 1 mM salicylic acid (SA) or mechanical damage.

(A–C) SA treatment strongly induces *PR1* expression but not *VSP1* compared with mock controls at 24 h. (A) Representative confocal image showing induced *PR1* expression but not *VSP1*. (B-C)

Quantification of *PR1* and *VSP1* expression after SA treatment. Images are maximum projection of confocal z-stacks. Scale bar, 100 μm.

(D–F) Mechanical damage induces *VSP1* expression but not *PR1* at 24 h. (D) Representative confocal image showing induced *VSP1* expression but not *PR1*. The dotted line indicates the damaged region. Scale bar, 200 μm. (E-F) Quantification of *PR1* and *VSP1* expression after mechanical damage.

Total fluorescence intensity density per image was measured. Scatter plots show median ± SD ($n \geq 6$ images from at least 3 plants). $p < 0.0001$, two-tailed Student's t-test.

**Movie S1 (separate file).** Z-stack movie showing strong *FRK1* expression in first-layer cells directly contacting bacterial colonies at early infection stages (related to Fig. 2A). Movie of a Z stack showing brightfield, mVenus, mCherry and chlorophyll signals from the adaxial upper epidermis to the mesophyll layer.

**Movie S2 (separate file).** Z-stack movie showing *FRK1* expression in cells adjacent to bacterial colonies (related to Fig. 2B).

**Movie S3 (separate file).** Z-stack movie showing redistribution of *FRK1* expression to second-layer cells at later infection stages (related to Fig. 2C–D).
